# Pathogenicity, strain properties and interspecies transmission capacity of pure recombinant prion protein assemblies

**DOI:** 10.1101/2023.02.27.530231

**Authors:** Human Rezaei, Davy Martin, Laetitia Herzog, Fabienne Reine, Naima Aron, Angélique Igel, Hannah Klute, Stella Youssafi, Jean-Baptiste Moog, Pierre Sibille, Olivier Andréoletti, Joan Torrent, Vincent Béringue

## Abstract

The pathogenicity of fibrillar assemblies derived from bacterially expressed recombinant prion protein (rPrP) has been key to the demonstration that prions are infectious proteins responsible for human and animal transmissible spongiform encephalopathies. Yet, their use in identifying which structural PrP features are important for prion biology, including strain properties and capacity to transmit between species, has been hampered by their limited transmissibility de novo. We report the generation of prions with distinct biological characteristics from rPrP assemblies differing only in their primary structure (hamster, mouse and human amino acid sequence). These rPrP assemblies were transmissible to transgenic mice expressing hamster PrP, causing a clinical disease at full attack rate, brain deposition of pathological prion protein PrP^Sc^ and spongiform degeneration. Their adaptation process on serial sub-passaging seemed to depend, as for genuine prions, on the presence of a species/transmission barrier, due notably to PrP sequence mismatch. Remarkably, one of the strains obtained is an unprecedented shortened prion, lacking the 90-140 amino-acid region which is believed to be key to infectivity and structural stability of disease-associated PrP assemblies. Finally, we provide evidence that rPrP prionogenicity lies in the structural organization and/or heterogeneity of the rPrP assemblies. These preparations of rPrP offer unprecedented opportunities for meaningful studies correlating the dynamicity and structures of PrP^Sc^ assemblies to prion pathobiology.

**Author summary:** Prions are infectious proteins, causing rapidly progressive neurodegenerative diseases in animals and humans. They are formed from the assisted-refolding and aggregation of the host-encoded prion protein (PrP). During the propagation of the disease, pathological PrP forces normal PrP to adopt its own conformation by a self-templating process. In infected host, different pathological structures of PrP or strains are found, causing diseases with specific biological phenotypes. Prions can also transmit between species. This capacity is limited by a species barrier, which critically depends on the infecting strain and PrP primary structure. How strain biological information is encoded in PrP structural fold remains unknown. We describe here the generation of different *bona fide* prion strains with markedly distinct adaptation capacities by transmission of refolded assemblies derived from bacterially-derived recombinant PrP (rPrP) of different species. We provide evidence that pathogenicity lies in the structural organization and/or heterogeneity of rPrP assemblies. Pathological PrP from one of the generated strains exhibited unique molecular features, including absence of domains that are thought to be key to prion infectivity, according to most recent ultrastructural studies. Our findings provide new insights for generating prion infectious material and resolving mechanisms of infectivity acquisition during PrP conversion process.

## Introduction

The mature form of the cellular prion protein (PrP^C^) is a ∼210 amino acids long, monomeric, GPI-anchored membrane glycoprotein with high primary and tertiary structure identities across mammals. PrP^C^ N-terminus is disordered; it contains tandem metal binding octarepeats and a hydrophobic region with a central polybasic domain. The C-terminus is folded and composed of three α-helices and a short antiparallel β-sheet [1, 2]. Mammalian prions are abnormally folded conformations (PrP^Sc^) of PrP^C^ with infectious properties. Misfolding of PrP^C^ into PrP^Sc^ involves enrichment in β-sheet content and assembling into polydisperse amyloidogenic assemblies [3]. PrP^Sc^ self-replicates by templating the conversion and polymerization of PrP^C^, causing inexorably fatal neurodegenerative diseases such as Creutzfeldt-Jakob disease (CJD) in humans, scrapie in sheep and goats, bovine spongiform encephalopathy (BSE) in cattle and chronic wasting disease in cervids [4].

Multiple prion strains are identified in the same host population. Strain properties, including incubation times, clinical signs and brain distribution of PrP^Sc^ deposition and vacuolar lesions are enciphered within structurally distinct PrP^Sc^ conformers [5]. Prion capacity to cross the species barrier varies among strains. At the molecular level, this reflects the capacity of the infecting PrP^Sc^ conformers to interact with PrP^C^ from a new host, i.e., with a primary structural mismatch [5, 6]. It is admitted that if these conformers lie within the portfolio of conformations host PrP^C^ can adopt in the PrP^Sc^ state, there will be little or no species barrier, and experimentally, propagation will occur at high attack rate, with minimal reduction of incubation periods and limited evolution of the strain phenotype on serial passage in the new host. If not, the species barrier will be high, with low attack rate and long incubation period on primary passage in the new host; adaptation will usually occur on serial transmission, sometimes abruptly, a phenomenon referred to as prion ‘mutation’ [7]. The main theoretical model describing prion cross-species adaptation is the “conformational selection” model [6, 8]. Based on the best replication selection [9], it considers the existence of structurally heterogenous PrP^Sc^ assemblies within a prion strain or isolate. The transmission barrier will favor the selection of the best replicator either because it pre-exists as an integral part of what defines a prion strain or because it is generated *de novo*, for instance by complementation between structurally distinct PrP^Sc^ assemblies [10]. A “deformed templating” variant of this model has been developed to account for the *in vivo* evolutionary process of certain recombinant prions [11]. The basic assumption of this variant is the absence of initial PrP assemblies heterogeneity which is compensated by creation of heterogeneity on initial passages in the new host, based on the templating interface misfit between PrP^Sc^ and host PrP^C^. Altered PrP^Sc^ conformers may be initially generated and outcompeted by the best replicant on serial adaptation.

While a global consensus on the *bona fide* infectivity of bacterially-derived recombinant PrP (rPrP) refolded *in vitro* into amyloid fibrils [12] has emerged, minimalistic (i.e., without co-factor(s) or auxiliary molecule addition) amyloid preparations often trigger incomplete attack rates, with long incubation periods or asymptomatic disease on initial passages to susceptible hosts [13–18]. In the deformed templating model, this limited transmissibility was interpretated as limited conversion capacity of PrP^C^ by rPrP fibrils due to absence of glycans and/or GPI. Others suggested low proportion of infectious conformers in the fibrillar samples, and/or cofactor(s) requirement, notably in the mouse brain, assisting the conversion or conferring infectivity. Indeed, co-incubation of rPrP with brain homogenate [11, 19] or specific auxiliary molecules, and/or extractive PrP^Sc^-seeded conversion of rPrP followed by amplification by chemically defined processes based on protein misfolding cyclic amplification (PMCA) [18, 20–25] allowed generating synthetic prions with significantly higher levels of infectivity. Given the difficulties to generate highly transmissible synthetic prions, the capacity of rPrP amyloid fibrils to generate different strains has rarely been addressed [15].

Here, we show that rPrP assemblies from diverse mammalian PrP primary sequences, formed spontaneously by partial unfolding in a minimalistic manner, are readily transmissible to hamster PrP transgenic mice and generate different strains. As for natural prions, the resulting strains readily replicate or need to adapt by crossing a species/transmission barrier. Structurally, these prions are composed of PrP^Sc^ conformers of low size, alone or in co-occurrence with ‘classical’ PrP^Sc^ conformers. Surprisingly, these low-size conformers are endogenously N-terminally truncated to an unusual degree of magnitude as they lack the 90-140 region which is believed to be key to prion infectivity.

## Results

### Structural heterogeneity of recombinant PrP assemblies

Recombinant PrP assemblies from hamster (rPrP^Ha^), human (rPrP^Hu^) and mouse (rPrP^Mo^) primary sequence were produced as described previously (**S1 Fig**, [26]). The assemblies were characterized for their size distribution, mean average molecular weight and morphology. As shown in **Fig 1A**, the dynamic and static light scattering analysis of the assemblies from the three species revealed a major hydrodynamic size distribution between 100 and 150 nm. The ultra-structure and morphology analyses of the assemblies by atomic force microscopy (AFM) using a 10Å nominal cantilever revealed an overall fibrillar shape (**Fig 1B**). However, numerous spherical objects were observed in every preparation. These spherical objects were either staked on the fibrils as protrusions or at vicinity or as isolated objects. The mean average Z-sizes of these spherical objects, as estimated from rPrP^Ha^, were around 25 Å and 40 Å (**Fig 1C**) whereas that of monomeric recombinant PrP is ∼12 Å. There was thus a constitutional heterogeneity within the generated rPrP assemblies used for the bioassay, with coexistence of at least two species. The term assembly rather than amyloid fibril will then be used to describe them.

**Figure 1.**
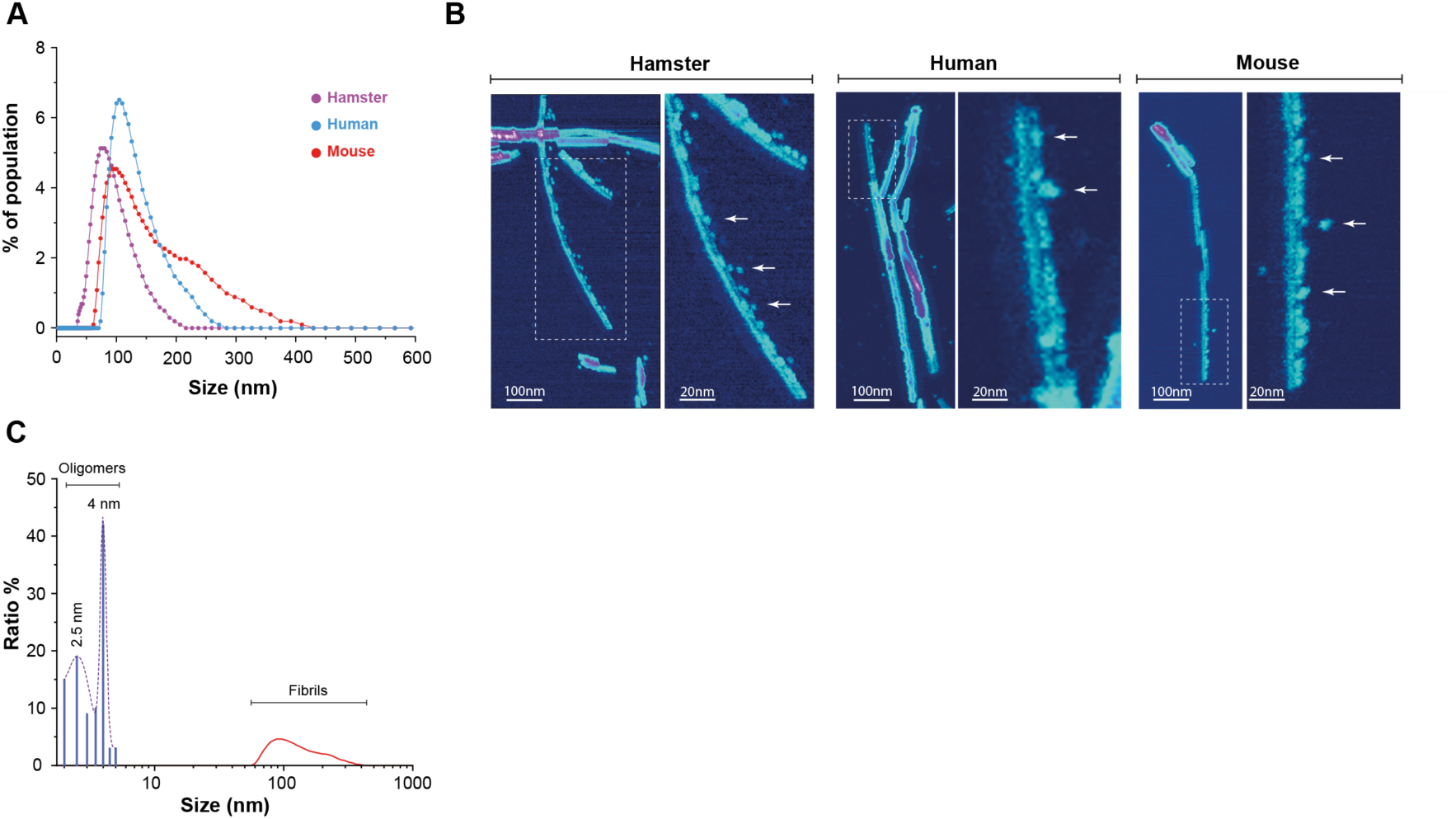
Quaternary structure characterization of mouse, hamster and human recombinant PrP assemblies. (**A**) Size distribution of the three recombinant PrP assemblies estimated by dynamic light scattering (DLS). The mean average hydrodynamic radius ranged between 100-150 nm. (**B**) Representative AFM images of spontaneously generated assemblies from hamster, human and mouse rPrP. Note the morphological heterogeneity for the three species with presence of fibrillar and spherical (arrows) assemblies. (**C**) Size characterization of the spherical assemblies estimated for hamster preparation. It indicates a typical 2.5nm and 4 nm population.

### Recombinant PrP assemblies are infectious

We inoculated 10 µg of hamster, mouse and human rPrP assemblies by intracerebral route to groups of 9-10 hamster PrP mice (tg7 line, [27]), overexpressing ∼4-fold brain PrP^C^ compared to hamster (**S2 Fig A-B**). As negative controls, tg7 mice were inoculated with rPrP monomers (10 per group; 10 µg). While none of the mice inoculated with the monomers developed any signs of disease up to >700 days post-inoculation (**S3 Table**), the mice inoculated with the assemblies developed a symptomatic neurological disease that was transmissible on subpassage, as summarized in **Table 1**.

**Table 1.** Serial transmission of hamster, mouse and human rPrP assemblies by intracerebral route in tg7 mice.

| rPrP assemblies | Mean survival in days ± SEM (n/n<sub>0</sub>)<sup>a</sup> | | |
| --- | --- | --- | --- |
|  | Hamster | Mouse | Human |
| 1<sup>st</sup> passage | 282 ± 17 (9/9) | 346 ± 10 (10/10) | 476 ± 11 (8/9) |
| 2<sup>nd</sup> passage | 273 ± 12 (8/8) | 84 ± 1 (8/8) | 260 ± 16 (8/8) |
| 3<sup>rd</sup> passage | 244 ± 13 (8/8) | 84 ± 2 (7/7) | 215 ± 7 (6/6) |
| 4<sup>th</sup> passage | 238 ± 15 (5/5) | 80 ± 2 (6/6) | 219 ± 7 (8/8) |

<sup>a</sup>n/n<sub>0</sub>: number of mice with neurological disease and positive for PrP<sup>res</sup> in the brain by immunoblotting/number of inoculated mice

On primary passage, a 100% and ∼90% attack rate was obtained with hamster/mouse rPrP assemblies and human rPrP assemblies, respectively. The mean survival times established at ∼280 days (rPrP^Ha^), ∼345 days (rPrP^Mo^) and ∼480 days (rPrP^Hu^). The clinical phase was protracted for the three groups; it lasted ∼4 weeks for rPrP^Mo^ assemblies and 4-6 weeks for rPrP^Ha^ and rPrP^Hu^ assemblies. By comparison, tg7-passaged hamster prions such as 263K, Sc237, 139H or ME7-H induce a short clinical phase lasting a few days [3, 10, 28]. Waddling gait, weight loss, kyphosis and a progressive paralysis affecting essentially the hindlimb, dominated in the three groups. Intense scraping and pruritus were noticed in tg7 mice inoculated with rPrP^Ha^ and rPrP^Hu^ assemblies.

Subpassages were performed with brains from individual, sick mice to investigate a potential transmission barrier or a low infectivity issue. With rPrP^Ha^ assemblies, the mean survival time on second passage was close to that seen on primary passage. It decreased modestly over further passaging to reach ∼240 days at the 4^th^ passage. With rPrP^Mo^ assemblies, the mean survival time abruptly decreased on second passage and stabilized at ∼80 days. With rPrP^Hu^ assemblies, the mean survival times decreased to attain ∼220 days at the 3^rd^ passage where it stabilized. Except for rPrP^Mo^ assemblies, the clinical phase remained prolonged on subpassaging for rPrP^Ha^ and rPrP^Hu^ assemblies.

Collectively, these data indicate that the three preparations of rPrP assemblies induced a *bona fide* transmissible disease in tg7 mice. The incubation time evolution over serial passaging suggests that at least two different prion strains emerged from these transmissions, one derived from mouse rPrP assemblies, the other one derived from hamster/human assemblies. To be noted, batches of rPrP^Ha^ and rPrP^Mo^ assemblies generated independently four years later were also fully pathogenic in tg7mice, with incubation times in the range of those initially observed (**S4 Fig A**). This observation indicates that the processes allowing the conversion of monomeric rPrP into infectious assemblies are deterministically governed.

### PrP^res^ electrophoretic signatures differ after infection with recombinant PrP assemblies

To confirm the occurrence of a *bona fide* prion disease after inoculation of rPrP assemblies and to characterize the strain phenotype obtained, we analyzed by immunoblot the brains of the diseased mice for the presence of PK-resistant PrP^Sc^ (PrP^res^) using the Sha31 anti-PrP antibody (central epitope, residues 145-152 [29]). All the brains accumulated detectable levels of PrP^res^ from the primary passage onwards. The electrophoretic profiles were unusual as compared to prototypical Sc237 PrP^res^ and variable among the three rPrP species. After infection with rPrP^Ha^ assemblies, PrP^res^ was characterized by the presence of three bands from ∼15 to ∼23 kDa (**Fig 2A, top panel**). This banding pattern was resolved in one band migrating at ∼15 kDa (**Fig 2A, middle panel**) after deglycosylation with PNGase, suggesting that the bands were variably glycosylated as Sc237 PrP^res^, but ∼5 kDa shorter in size. The unique presence of this low molecular weight (LMW) pattern was observed over 4 passages. Mice infected with the second batch of rPrP^Ha^ assemblies showed a similar LMW PrP^res^ banding pattern in the brain (**S4 Fig B**), indicating an invariance in the banding pattern as the function of the batch.

**Figure 2.**
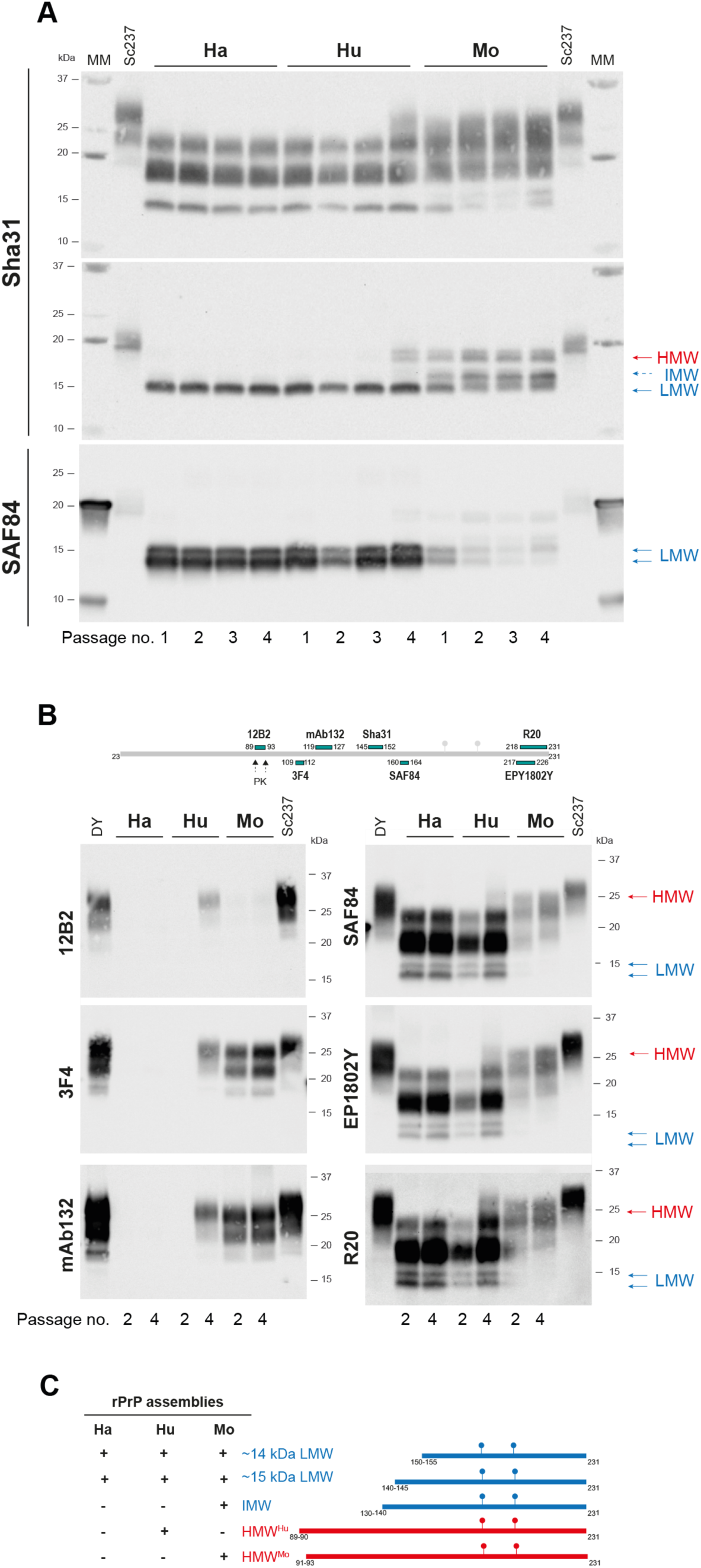
Electrophoretic profiles of PrP^res^ detected in the brains of tg7 mice inoculated with recombinant PrP assemblies. (**A**) Electrophoretic pattern of PrP^res^ in the brains of tg7 mice on serial passage of rPrP assemblies from hamster (Ha), human (Hu) and mouse (Mo) species. The pattern is shown before (top panel) and after (middle and bottom panels) deglycosylation by PNGase. Immunoblots were probed with Sha31 and SAF84 anti-PrP monoclonal antibodies, as indicated. As control, the electrophoretic pattern of Sc237 in tg7 mice is shown. The passage number (no.) is indicated. HMW, IMW and LMW PrP^res^ species are indicated by an arrow. MM: molecular mass markers. (**B**) Epitope mapping of PrP^res^ at 2^nd^ and 4^th^ passage (as indicated). The antibodies used to probe the blots and their epitopes on PrP are indicated in the schematic diagram. As controls, the electrophoretic pattern of PrP^res^ from tg7-passaged Sc237 and DY prions are shown. (**C**) Inventory of the PrP^res^ fragments found in prions of hamster (Ha), human (Hu) and mouse (Mo) rPrP origin (4^th^ passage in tg7 mice).

Mice inoculated with rPrP^Hu^ assemblies exhibited the same LMW PrP^res^ banding pattern over 3 passages, (**Fig 2A**). However, at the 4^th^ passage, a second PrP^res^ product of higher molecular weight (HMW) was detected, which, after deglycosylation, migrated slightly faster than Sc237 PrP^res^ (**Fig 2A**).

After infection with rPrP^Mo^ assemblies, PrP^res^ was characterized by the presence of both LMW and HMW products in variable proportions, from the primary passage onwards (**Figure 2A**). After deglycosylation, a third product of intermediate molecular weight (IMW), migrating at ∼16 kDa, was detected (**Fig 2A**). The same PrP^Sc^ signature heterogeneity was conserved with the second batch of rPrP^Mo^ assemblies (**S4 Fig B**).

To further resolve the assemblies-specific PrP^res^ banding patterns, we performed an epitope mapping with monoclonal antibodies against PrP, with epitopes spanning the PrP amino acid sequence from the PK-cleavage sites up to the C-terminus (**Fig 2B**). This was done on brain samples at the 2^nd^ and 4^th^ passage. The LMW PrP^res^ pattern was not detected in any of the inoculated mice when immunoblots were probed with 12B2 (PrP epitope 89-93 [30]), 3F4 (PrP epitope at residues 109 to 112 [31]) or mAb 132 (PrP epitope at residues 119-127 [18]) anti-PrP antibodies (**Fig 2B, left column**), whatever the passage analyzed and the rPrP assembly inoculated. The SAF 84 antibody which has an epitope in the central region of PrP after Sha31 (PrP epitope at residues 160 to 164 [32]) and antibodies with an epitope in the C-terminal part of PrP (EPY1802Y (PrP epitope 217-226) and R20 (PrP epitope 218-231) [18]) recognized LMW PrP^res^ from tg7 mice inoculated with hamster, human and mouse rPrP assemblies (**Fig 2B, right column**). With these three antibodies, but not with Sha31, a second LMW band migrating ∼1 kDa faster was detected (**Fig 2B, right column and 2A, bottom panel**). By considering the average weight of one amino acid, the size of the GPI anchor, the size calculated with our molecular mass markers and Sha31/SAF84 epitope, we estimated that the ∼15 kDa LMW PrP^res^ would have a sequence starting from amino acid 140-145 until the C-terminal 231 amino acid. The ∼14 kDa LMW PrP^res^ detected with SAF84 and the C-terminal anti-PrP antibodies would have a sequence starting from amino acid 150-155.

The HMW PrP^res^ pattern seen at the 4^th^ passage of rPrP^Hu^ assemblies (HMW^Hu^) and with rPrP^Mo^ assemblies (HMW^Mo^) was detected with all the anti-PrP antibodies tested (**Fig 2B**). However, with 12B2 anti-PrP antibody, the HMW^Mo^ PrP^res^ signal almost disappeared compared to HMW^Hu^ or tg7-passaged Sc237 and DY. In addition, probing with mAb132 and 3F4 anti-PrP antibodies showed that HMW^Mo^ PrP^res^ migrated slightly faster than HMW^Hu^ PrP^res^. This suggested that HMW^Mo^ PrP^res^ was slightly more N-terminally truncated than HMW^Hu^ PrP^res^ beyond PK-cleavage site at position 89-90.

The IMW PrP^res^ banding pattern found after mouse rPrP assemblies inoculation (IMW^Mo^) was not positive with 12B2, 3F4 and mAb 132 anti-PrP antibodies (**Fig 2**). This fragment was thus truncated between amino acid position 130 and 140.

Collectively, classification based on electrophoretic signatures of the PK-resistant PrP^Sc^ core over passaging allows further distinguishing between prions of hamster, human and mouse rPrP origin (**Fig 2C**). We thus isolate three different prion strains in tg7 mice. rPrP^Ha^-derived prions were solely composed of two LMW fragments that were N-terminally truncated up to residues 140-145 and 150-155. rPrP^Hu^-derived prions show additional presence of HMW^Hu^ PrP^res^ starting at position ∼90 from the 4^th^ passage onwards. rPrP^Mo^-derived prions show additional presence of HMW^Mo^ and IMW^Mo^ PrP^res^ species starting at position ∼93 and 130-140, respectively. The coexistence of several proteolytic fragments specifically in the case of human and mouse rPrP assemblies indicates a high degree of structural heterogeneity of their respective PrP^res^ assemblies.

### LMW PrP^Sc^ is mostly endogenously truncated

To determine if LMW PrP^res^ results from the partial digestion of full-length PrP^Sc^ by PK, we digested brain homogenates from mice inoculated with hamster, human and mouse rPrP assemblies (2^nd^ and 4^th^ passage) with thermolysin instead of PK. For multiple prion strain/host PrP combinations, including hamster PrP, limited digestion with thermolysin is known to completely degrade PrP^C^ while leaving >80 % of PrP^Sc^ as full-length form [33–35]. Mock-infected brain homogenates from tg7 mice were used as control for efficient digestion of PrP^C^ by thermolysin. As positive control, Sc237-infected brain homogenates were used. After digestion and SDS-PAGE electrophoresis, the immunoblots were revealed with anti-PrP antibodies with epitopes in the N-terminal domain (SAF34, human PrP epitope at octarepeats residues 59 to 89 [32]) or C-terminal domain of PrP (Sha31 and SAF84) to check for full-length PrP^Sc^ and full-length + truncated PrP^Sc^, respectively. The results are summarized in **Fig 3A**. Whatever the antibody used for immunoblotting, no signal was detected in uninfected brain homogenates, suggesting complete enzymatic digestion of PrP^C^. PrP^Sc^ from Sc237 prions was detected mostly as full-length PrP^Sc^ as inferred from the presence of SAF34-positive signal and the faint, extended pattern seen with both Sha31 and SAF84 antibodies. Thermolysin-digested PrP^Sc^ from rPrP^Ha^-derived prions at 2^nd^ and 4^th^ passage and from rPrP^Hu^-derived prions at 2^nd^ passage were detected at low levels with SAF34 while strongly positive, as a LMW truncated form with Sha31 or as two LMW truncated forms with SAF84. Thermolysin-digested PrP^Sc^ from rPrP^Hu^-derived prions at 4^th^ passage and from rPrP^Mo^-derived prions at 2^nd^ and 4^th^ passage yielded a stronger SAF34 positive signal, thereby suggesting enriched presence of full-length PrP^Sc^.

**Figure 3.**
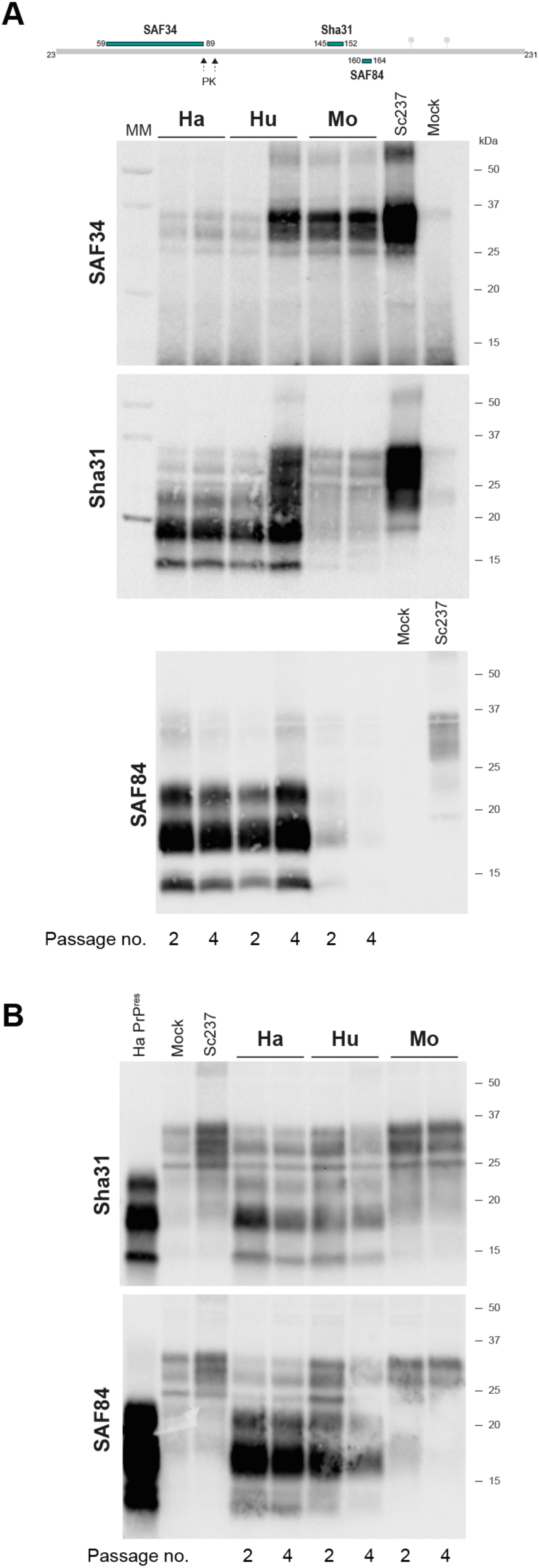
Thermolysin-resistance and insolubility of PrP^Sc^ present in the brains of tg7 mice inoculated with recombinant PrP assemblies. Electrophoretic pattern of (**A**) thermolysin-resistant PrP^Sc^, (**B**) insoluble PrP^Sc^ in the brains of tg7 mice on serial passage of rPrP assemblies from hamster (Ha), human (Hu) and mouse (Mo) species. The analysis was performed at the 2^nd^ and 4^th^ passage, as indicated. As controls, thermolysin digested or ultracentrifuged brain homogenates from Sc237 infected or mock-inoculated tg7 mice are shown. In panel (**B**), the electrophoretic pattern of PrP^res^ in the brain of tg7 mice on serial passage of rPrP^Ha^ assemblies is shown (labeled HaPrP^res^). The antibodies used to probe the blots and their epitopes on PrP are indicated in the schematic diagram. MM: molecular mass markers.

Collectively, these data indicate that the LMW PrP^Sc^ species are mostly truncated prior to PK digestion. rPrP assemblies may convert full-length PrP^C^ in a PrP^Sc^ conformation that is either unusually sensitive *in vitro* to thermolysin or *in vivo* to endogenous proteolytic cleavage [36], yielding LMW species.

To distinguish between these two hypotheses, we analyzed PrP^Sc^ electrophoretic pattern in solubilized brain homogenates from mice inoculated with hamster, human and mouse rPrP assemblies (2^nd^ and 4^th^ passage) by sedimentation assay in the absence of any protease treatment. The immunoblots were revealed with anti-PrP antibodies Sha31 and SAF84 to check for full-length + truncated PrP^Sc^. The results are summarized in **Fig 3B**. A faint signal was detected in solubilized+pelleted uninfected brain homogenates, suggesting limited insolubility of PrP^C^. PrP^Sc^ from Sc237 prions was mostly detected as full-length form. Sedimentable PrP^Sc^ species from rPrP^Ha^-derived prions and from rPrP^Hu^-derived prions were mostly detected as LMW truncated forms with Sha31 and SAF84 antibodies. Monoglycosylated dominant full-length PrP^Sc^ (i.e., different from the signal obtained with uninfected brain) was detected as minor component with these antibodies. PrP^Sc^ from rPrP^Mo^-derived prions was found predominantly as full-length PrP^Sc^. These data thus indicate that LMW PrP^Sc^ is mostly endogenously truncated. This suggests that rPrP assemblies convert either full-length PrP^C^ in a PrP^Sc^ conformation sensitive to endogenous proteolytic cleavage or a truncated form of PrP^C^ that yields directly, or after cleavage, LMW species.

### Neuropathology of mice infected with recombinant PrP assemblies

We next studied by histoblotting the strain-specified [37] neuroanatomical distribution of PrP^res^. **Fig 4** synthetized the results obtained at the 4^th^ passage, i.e., when the incubation times were stabilized. To detect LMW PrP^res^ species, immunoblotting was performed with SAF84 anti-PrP antibody. The 3F4 antibody was used additionally with rPrP^Mo^-derived prions to detect HMW^Mo^ PrP^res^ (note that the levels HMW^Hu^ PrP^res^ were too low to be specifically detected by histoblotting with 3F4). Mice infected with rPrP^Ha^- and rPrP^Hu^-derived prions exhibited similar PrP^res^ deposition pattern. There was a widespread deposition of PrP^res^ aggregates in several brain regions, including the cerebral cortex, the thalamus, and the brain stem. Several white matter tracts scored strongly positive, including the corpus callosum, the external capsule, the cingulum, the optic and olfactory tracts. LMW PrP^res^ deposition in these tracts was more visible at 3^rd^ passage as the deposition throughout the brain was less widespread (**S5 Fig**). After infection with rPrP^Mo^-derived prions, SAF84 detected low levels of PrP^res^ preferentially in these white matter tracts (**Fig 4 and S5**). As this antibody detects preferentially LMW PrP^res^, this suggests overlapping deposition of LMW PrP^res^ forms after inoculation with the three rPrP-derived prion strains. With 3F4 antibody, mice infected with rPrP^Mo^-derived prions exhibited a HMW PrP^res^ deposition pattern that broadly overlapped that of LMW PrP^res^ with respect to the widespread deposition. In addition to the aforementioned brain regions, PrP^res^ deposited in the hippocampus and the caudate putamen (**Fig 4**). Altogether there were many regions of co-deposition of HMW PrP^res^ and LMW PrP^res^, suggesting at the resolution of the histoblots, absence of specific (truncated) PrP^C^ substrate converted into LMW PrP^res^.

**Figure 4.**
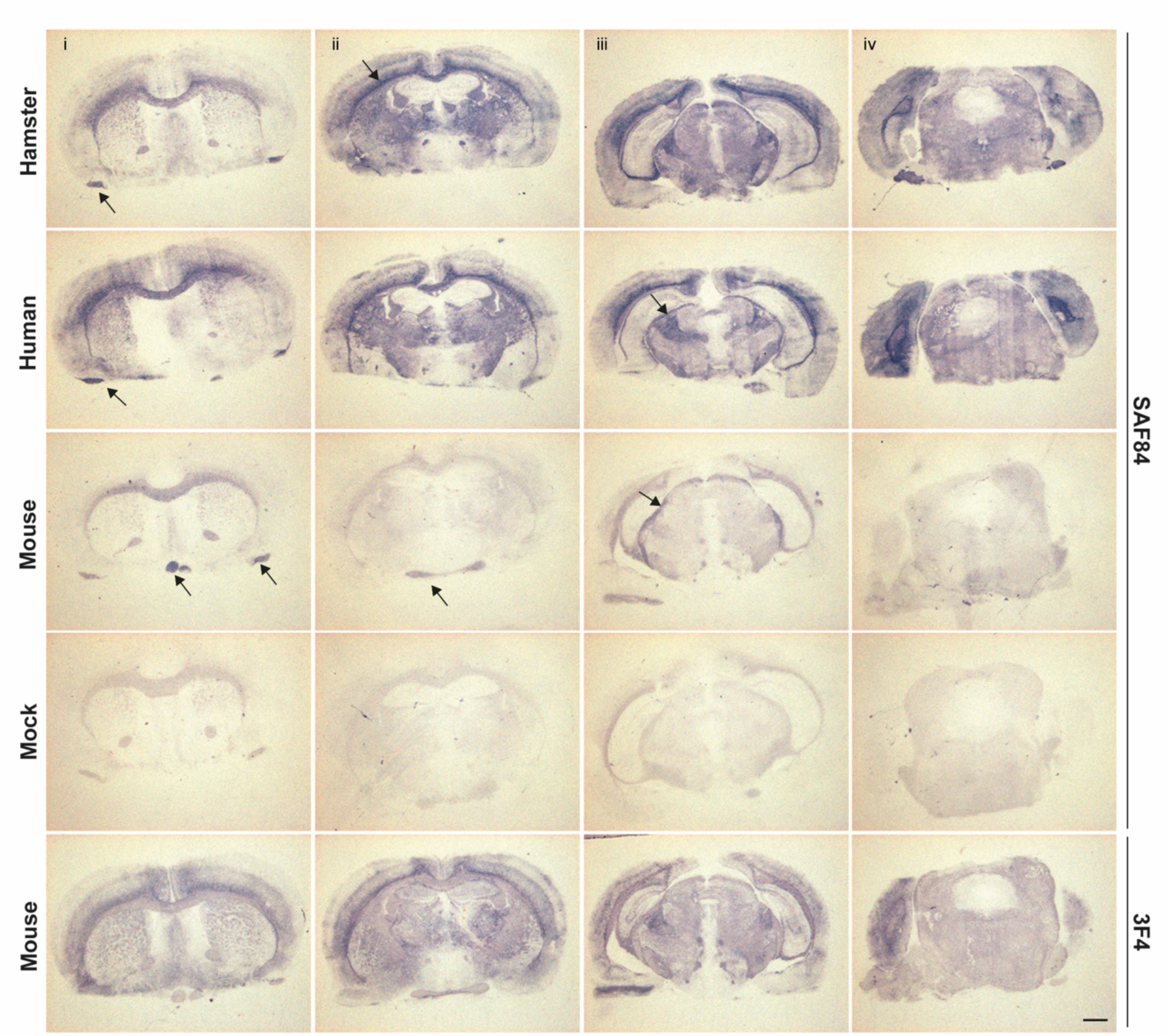
Neuropathological pattern of PrP^res^ deposition in tg7 mice inoculated with recombinant PrP assemblies. Representative histoblots of rostro-caudal transversal brain sections are shown at the 4^th^ passage, after challenge with rPrP assemblies from hamster (Ha), human (Hu) and mouse (Mo) species. Analyses were performed at the level of the septum (i), hippocampus (ii), midbrain (iii) and brainstem (iv). As control, histoblots from mock-infected tg7 brain are shown. Histoblots were probed with 3F4 and SAF84 anti-PrP monoclonal antibodies, as indicated. Arrows indicate PrP^res^ deposition in white matter tracts. Scale bar, 1mm.

To complete this analysis, we established by histological examination strain-specific [38] vacuolar lesion profiles (**Fig 5A**). Spongiform degeneration was more severe and widespread after inoculation with hamster, human and mouse rPrP assemblies as compared to Sc237-inoculated tg7 mice. The spatial variability of the brain lesion profile was limited on subpassaging for the three strains. Mice inoculated with rPrP^Ha^ and rPrP^Hu^ assemblies exhibited relatively similar lesion profile and the intensity of vacuolation increased on subpassaging. The lesion profiles were different from that seen in mouse infected with rPrP^Mo^ assemblies. With rPrP^Ha^- and rPrP^Hu^-derived prions, there was intense areas of vacuolation in the cortex and hippocampus, which was accompanied by visible neuronal loss (**Fig 5A-B**).

**Figure 5.**
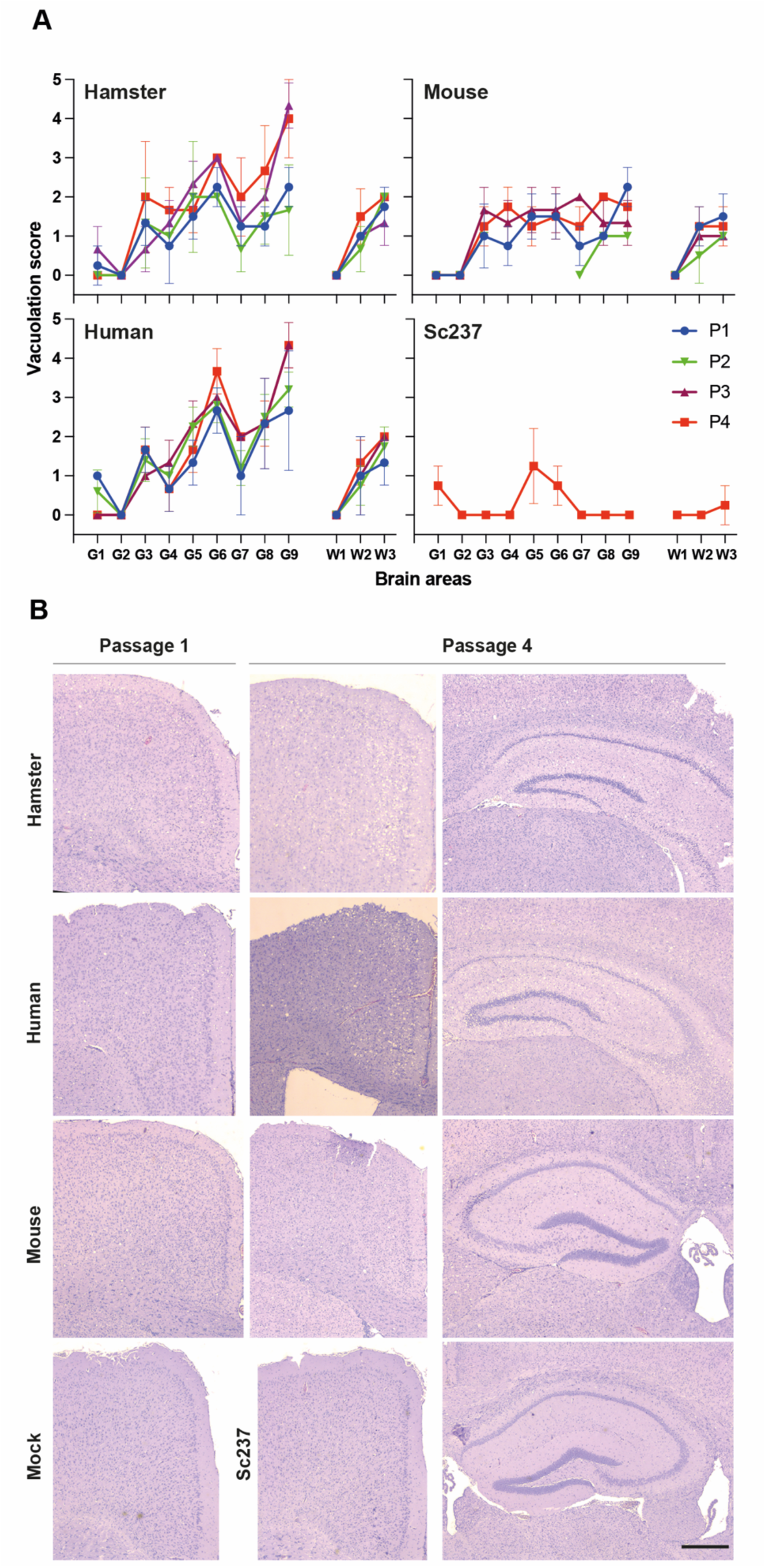
Spongiform degeneration in tg7 mice inoculated with recombinant PrP assemblies. (**A**) Standard vacuolar lesion profiles in the brains of tg7 mice on serial passage (P) of rPrP assemblies from hamster, human and mouse species. Sc237 in tg7 mice was used as control. The vacuolation intensity was scored as means ± SD in standard gray (G1-G9) and white (W1-W3) matter areas. G1: dorsal medulla; G2: cerebellar cortex; G3: superior colliculus; G4: hypothalamus; G5: medial thalamus; G6: hippocampus; G7: septum; G8: medial cerebral cortex at the level of the thalamus; G9: medial cerebral cortex at the level of the septum; W1: cerebellar white matter; W2: white matter of the mesencephalic tegmentum; and W3: pyramidal tract. (**B)** Spongiform changes in tg7 mice at the first and fourth passage. Note the neuronal degeneration in the pyramidal cell layer of the hippocampus with rPrP^Ha^- and rPrP^Hu^-derived prions (scale bar: 400 µm).

Collectively, these data indicate that rPrP assemblies triggered prion-specific neuropathological stigmata. They further differentiate synthetic rPrP^Mo^-derived prions from rPrP^Ha^- and rPrP^Hu^-derived prions. Finally, they lend support to the view that LMW PrP^Sc^ species that are dominant in rPrP^Ha^-derived prions are fully pathogenic *per se*.

### PrP misfolding pathway determines synthetic prion genesis

To finally determine the importance of the misfolding PrP pathway in the generation of infectious rPrP assemblies, we used a different methodology. Instead of using partial destabilizing conditions using chaotropes to product rPrP fibrillar assemblies, we performed a thermal treatment. Thermal treatment of hamster, human and mouse monomeric rPrP at acidic pH mainly generates ß-sheet-rich oligomeric assemblies that bind thioflavin T [39]. However, these oligomeric objects are unstable at pH 7.0 and depolymerize into monomeric PrP, thus necessitating the use of rPrP from another mammalian species to address the PrP unfolding/misfolding pathway-infectivity relationship. Thermal treatment of sheep rPrP (V_136_R_154_Q_171_ allele) generates high molecular weight oligomers, heterogenous in size (> 36 PrP-mers), highly stable with high ß-sheet content, thioflavin T-positive [40, 41] and able to condensate into molecular assemblies presenting a nodular fibrils shape (**Fig 6A**). The same monomeric sheep rPrP by chaotropic treatment gave rise to assemblies morphologically similar to those obtained with hamster, human and mouse rPrP, with presence of fibrillar and spherical structures (**Fig 6A**).

**Figure 6.**
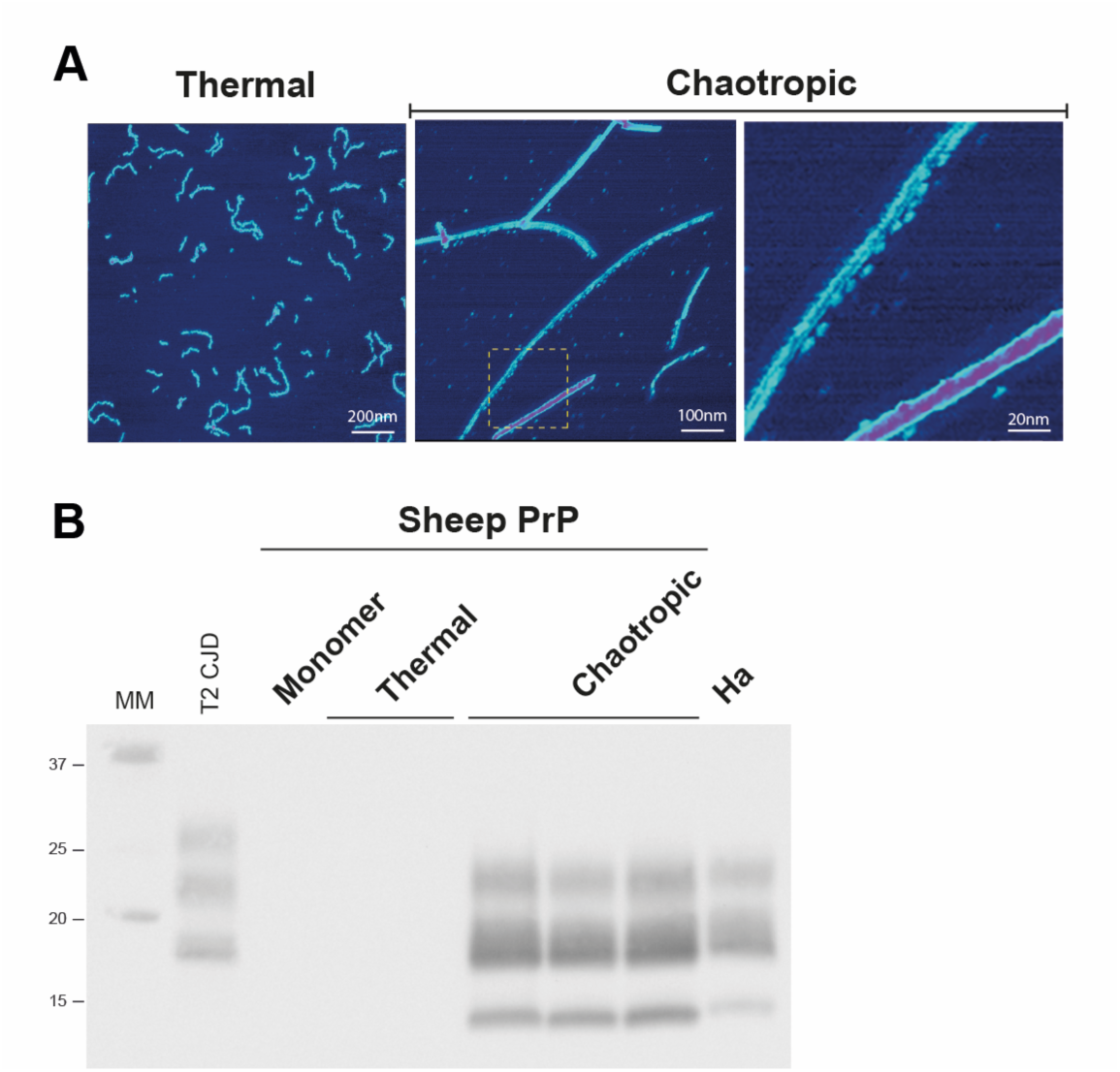
Morphology and transmission properties of recombinant PrP assemblies generated by thermal versus chaotropic treatment. (**A**) Representative AFM images of sheep rPrP assemblies spontaneously generated by thermal or chaotropic treatment. The assemblies were deposited on mica and imaged with high-resolution AFM tips. (**B**) Detection and electrophoretic pattern of PrP^res^ in the brains of tg7 mice on primary passage of sheep rPrP monomers and assemblies generated by thermal versus chaotropic treatment. As controls, LMW hamster PrP^res^ (Ha) and T2 CJD PrP^res^ are shown. Immunoblots were probed with Sha31 anti-PrP monoclonal antibody. MM: molecular mass markers.

We thus compared the transmission properties of sheep rPrP assemblies generated by thermal versus chaotropic treatment. Those were inoculated by intracerebral route to groups of 8-11 tg7 mice, at the 0.5 mg/mL final concentration. As control, we inoculated monomers at the same concentration. None of the mice inoculated with the monomers or with the assemblies obtained by thermal treatment developed any signs of disease nor accumulated PrP^res^ in their brains (**Table 2, Fig 6B**). In contrast, tg7 mice inoculated with sheep rPrP assemblies obtained by chaotropic treatment developed a clinical disease at 100% attack rate (**Table 2**).

**Table 2.** Primary transmission of sheep rPrP monomers and assemblies obtained by thermal or chaotropic treatment by intracerebral route in tg7 mice.

| rPrP | Survival<sup>a</sup> time in days (n/n<sub>0</sub>)<sup>b</sup> |  |  |
| --- | --- | --- | --- |
|  | Monomers | Thermal | Chaotropic |
| 1<sup>st</sup> passage | 312-636 (0/8) | 428-768 (0/8) | 326 ± 12 (11/11) |

<sup>a</sup>For the negative transmissions, the range of individual survival times is given. For the positive transmission, survival is expressed as mean ± SEM.
<sup>b</sup>n/n<sub>0</sub>: number of mice with neurological disease and positive for PrP<sup>res</sup> in the brain by immunoblotting/number of inoculated mice

PrP^res^ was detected in the brain of these mice and exhibited a LMW PrP^res^ profile (**Figure 6B**). Collectively, these data suggest that the spontaneous formation of infectious assemblies depends on the nature of perturbation of the native state and thus the unfolding/misfolding pathway.

## Discussion

There has been considerable literature on the infectivity of recombinant PrP assemblies. The limited transmissibility of crude preparations and the difficulty to reproduce certain experiments [42] has made the interpretation of certain analyses difficult, notably those designed to explore the biochemistry of the infectious species and the propagation/adaptation process in vivo. Here, we report the transmissibility, strain properties and evolutive/adaptative potential of minimalistic preparations of fibrillar assemblies from recombinant PrP with amino acid sequence from four different mammalian species. We reach two main conclusions from our studies: first, chaotropic treatment of monomeric rPrP from different species spontaneously generates distinct prion strains. As for ‘natural’ prions, rPrP assemblies ‘faithfully’ propagated in the infected host or required serial passaging to adapt in the homotypic PrP and heterotypic PrP context, respectively. Second, the resulting rPrP-derived prions were biochemically characterized by the presence of an endogenously generated, truncated C-terminal PrP^Sc^ fragment, suggesting that the beginning of the globular domain in PrP N-terminus is dispensable to PrP^Sc^ infectious core and thereby questioning the unicity of a common structural model thereof. We also discuss the potential importance of the structural heterogeneity of our rPrP preparations, notably the presence of non-fibrillar assemblies, in conferring infectivity to recombinant PrP.

### The species barrier primarily dictates rPrP assemblies transmission efficacy

Transgenic mice expressing hamster PrP were uniformly susceptible to hamster, human and mouse rPrP assemblies, according to the high attack rate, presence of stereotyped clinical symptoms, PrP^Sc^ detection and spongiform degeneration, from the primary passage onward. Primary passage of sheep rPrP assemblies in these mice also suggests efficient transmission capacity. Hamster, human and mouse rPrP assemblies encoded distinct prion strains, as based on strain typing analyses over 4 serial passages. With rPrP^Ha^ assemblies, full attack rate, modest evolution of the mean survival time over 4 passages (<15%) and stability of the biochemical and neuropathological phenotypes suggest efficient replication with minimal adaptation. It extends, with prions of recombinant origin, the importance of PrP sequence homology in abrogating prion transmission barriers [43–45]. Mouse and human rPrP assemblies induced disease at complete or near-complete attack rate on primary passage. The incubation times evolved substantially on subpassage, either abruptly to become very short with rPrP^Mo^ assemblies or at slower pace with rPrP^Hu^ assemblies. The PrP^res^ signature rPrP^Hu^-derived prions also evolved on serial passage. These transmission properties were paralleling those obtained on evolution of prions in the context of a species barrier or a transmission barrier, with slow [46, 47] or abrupt (‘mutational’) adaptation [7, 48]. These findings extend, with prions of recombinant origin, the importance of the strain type in crossing prion species barriers [49, 50]. Overall, the transmission efficacy of rPrP assemblies appears primarily driven by the potential presence of a species or transmission barrier.

### Relative infectivity of recPrP assemblies and comparison with natural prions species barrier

The magnitude of the transmission barrier obtained with rPrP assemblies in tg7 mice is consistent with that seen with natural strains as shown in **Fig 7**. Tg7, and more generally transgenic mouse models, do not behave differently than wild-type animals with respect to prion transmission barrier **(S2 Fig C** and ref. [51] for an overview). We recently suggested that during prion adaptation to a new host, two distinct subpopulations of PrP^Sc^ helped crossing the species barrier by structural complementation [10, 52]. The synergetic effect between these two subpopulations present in the inoculum of natural prion strains makes serial dilution on primary passage aberrantly decreases the transmission yield by several orders of magnitude in the presence of a transmission barrier [10]. The comparison of disease durations obtained with rPrP^Ha^ assemblies and hamster-adapted prions (**Fig 7**) does not provide information on their relative infectivity levels because disease duration is entangled with toxicity [53, 54]. Yet, in the case of rPrPHa it should be noted that disease duration shortening between 1^st^ and 2^nd^ passage is like other hamster-adapted strains, suggesting a stable level of infectivity over passaging and an absence of species barrier. By taking 263K/Sc237 prions as reference, a rapid estimation based on the amount of total PrP^Sc^ tends to indicate that the initial specific infectious titer of rPrPHa was roughly 20-fold less than for 263K/Sc237 prions (**S6 Text**).

**Figure 7.**
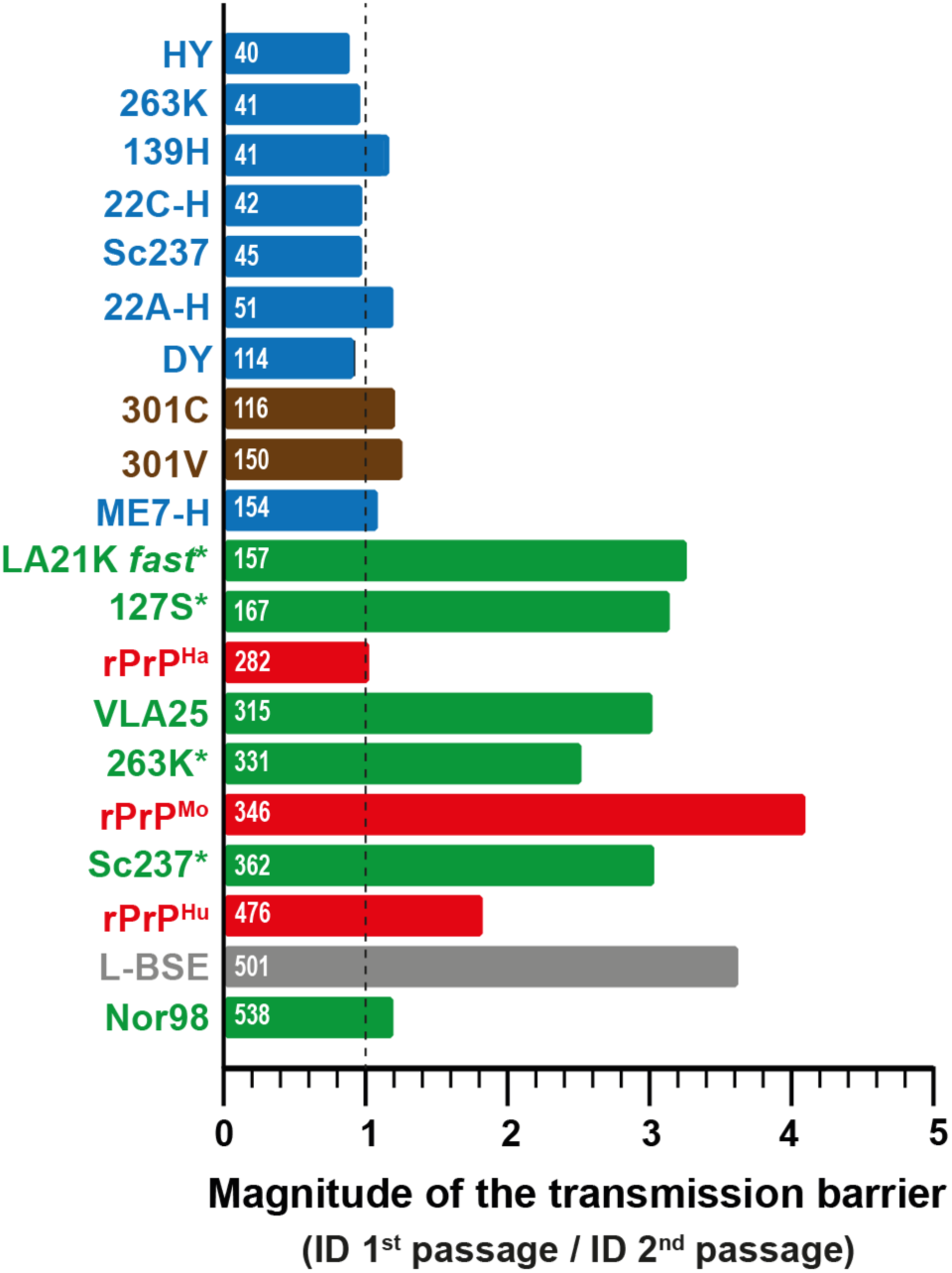
Comparison of the magnitude of the species/transmission barrier between recombinant PrP assemblies and natural prion strains on transmission to tg7 mice. The transmission properties of hamster-adapted prions (blue), mouse-adapted BSE prions (brown), ovine-adapted prions (tg338 mice (*)[10, 77] or sheep, green), bovine L-BSE prions (gray) and rPrP assemblies (red) are shown. The mean incubation time on primary passage are indicated in white color. The magnitude of the transmission barrier is calculated as the ratio of the mean incubation durations (ID) on first to second passage in tg7 mice [10, 50]. A value of 1 (dash line) suggests absence of transmission barrier.

The pathogenicity of our rPrP preparations contrast with previous studies which reported incomplete attack rates, long incubation periods or asymptomatic disease in bioindicator animals inoculated with similar concentration of rPrP fibrils made sometimes in quasi-similar conditions [14–17]. Intriguingly, inoculation of hamster rPrP fibrils to hamster or to a hamster PrP transgenic mouse line distinct from ours led, over 2 passages, to few asymptomatic animals accumulating a misfolded form of PrP^Sc^ in their brain, with LMW-like electrophoretic signature. This signature was termed ‘atypical’ PrP^res^. On serial subpassaging, atypical PrP^res^ gradually transformed into classical, Sc237-like PrP^res^, which eventually became the predominant species, correlating with the development of a clinical disease, at least in some animals. The “deformed-templating” hypothesis [11] was developed to explain these results. We show a fundamentally opposite situation with rPrP^Ha^ assemblies preparations and our hamster PrP transgenic mouse model, which is not different with respect to the transgene used and overall PrP overexpression levels ([55–57], **S2 Fig**). Furthermore, LMW PrP^res^ and ‘atypical’ PrP^res^ looked identical as based on electrophoretic mobilities and epitope mapping [16, 18]. The evolution of prions originating from rPrP^Hu^ and rPrP^Mo^ assemblies also led to a different outcome than reported for the deformed-templating experiments. With rPrP^Hu^ assemblies, only LMW PrP^res^ was detected over 3 passages. At the 4^th^ passage, HMW^Hu^ PrP^res^ species emerged, without any significant impact on the disease incubation time or neuropathology. With rPrP^Mo^ assemblies, both LMW and HMW^Mo^ PrP^Sc^ species were detected in the brains from the first passage onward; both species were detected independently of the disease evolution on subpassaging. Recombinant PrP assemblies thus behave or evolve molecularly like naturally occurring prions as emergence and/or co-propagation of distinct PrP^Sc^ conformations (associated sometimes with distinct strains) is far from unusual, particularly following interspecies transmissions [7, 45, 58–61].

### Recombinant PrP assemblies heterogeneity may facilitate transmission

There has been a systematic investigation of physico-chemical conditions to generate highly infectious mouse rPrP fibrils [13]. Yet, the authors used as readout for prion infectivity a scrapie cell assay that may not replicate slowly replicating rPrP-derived prions. Other studies on synthetic prions used fibrillar preparations purified to structural homogeneity or used preparations generated by in vitro self-amplification in the potential presence of different cofactors. This may have selected certain assemblies and/or modified the relative ratio of conformations [62], resulting in delayed or inefficient transmission. This is a common practice in soft matter biochemistry and amyloid structural biology to tune physico-chemical conditions to obtain a high degree of structural homogeneity. Why our rPrP assemblies preparations are readily infectious? A first hypothesis is that the physico-chemical conditions used to prepare rPrP assemblies may impact the amount of the replicative population. Accordingly, other rPrP structuration, obtained by thermal instead of chaotropic treatment, were not infectious on primary transmission to hamster PrP mice, despite high ß-sheet content and thioflavin T positivity. A second (non-exclusive) hypothesis could be the degree of heterogeneity and the presence of a “helper” assembly, which may confer enhanced transmissibility. We recently reported that the early steps of prion conversion intrinsically generated two sets of infectious/replicative PrP^Sc^ assemblies, a small oligomeric species of a size smaller than a pentamer and a larger species of a size higher than a MDa. These two species affect their own formation through a kinetical feedback [62] and their synergetic action is mandatory for prion adaptation and the passage of a species barrier [10]. The infectivity titer of our preparations could lay on the coexistence of spheric species with the fibrillar objects. Ultrastructural analyses by AFM of the assemblies preparations obtained by chaotropic treatment of the four mammalian rPrP revealed indeed a relatively high degree of heterogeneity, which was not observed with the assemblies generated by thermal treatment.

### An infectious N-terminally truncated mini-prion

The LMW PrP^res^ strain signature is biochemically distinguishable by the characteristic size of the fragments. It migrated from ∼14-15 to ∼22-23 kDa, depending on its glycosylation status. Epitope mapping indicates that LMW PrP^res^ is composed of 2 N-terminally truncated fragments starting around amino acids ∼140-145 and 150-155, instead of ∼90 for classical PrP^res^. Use of thermolysin as chemical probe and differential centrifugation further indicated that LMW PrP^res^ was mostly truncated in vivo. Whether this occurs through an endogenous degradative process post full-length PrP^C^ conversion or due to straight conversion of prionogenic PrP^C^ fragment(s) of shorter size remains to be determined [63, 64]. Whatever their origin, these shortened PrP^Sc^ assemblies may contain de novo infectivity and neurotoxic determinants. It thus raises the question of the respective contribution to neurodegeneration of these conformations versus classical PrP^Sc^ when both are co-propagated in the brain of prion-infected animals [29, 65] or humans [66].

Recently, we showed that the alpha-cleaved C1-fragments from PrP^C^ (truncation at amino acids 113/115, [67]) were convertible into prions in RK13 cells, when part of the α-helix H2 was deleted [68]. This result and the infectivity of LMW PrP^Sc^ fragments stand in apparent contradiction with several reports suggesting that PrP amino acids ∼90-100 to 155, notably the central polybasic domain with 4 lysines at aa ∼100-110, are an obligate domain of PrP^res^ infectious core [18, 69–71]. Maybe the most illustrative results are the transmission properties of shortened mouse rPrP fibrils (residues 23-144) corresponding to the Y145Stop PrP variant associated with the Gerstmann-Sträussler-Scheinker syndrome. They were infectious in mouse Tga20 mice, with minimal adaptation on serial passaging [22]. This would correspond to the domain that is absent in LMW PrP^res^. The fold allowing prion infectivity and strain structural determinant can thus be subject to major variations with respect to the PrP domains involved.

### Are LMW prions compatible with recent prion architectures?

Recent cryo-EM studies of purified PrP^res^ amyloid fibrils from hamster 263K and mouse RML or ME7 prion strains revealed a single- or paired-protofilament helical structure with a parallel-in-register intermolecular β-sheet (PIRIBS)-based core architecture [72–74]. Applying the hamster 263K-PIRIBS architecture to hamster LMW PrP^res^ would remove ∼35% of the β-strands (β1 to β4 444 in the N-terminus lobe), and in terms of structural elements, the zipper that seals residues 96-100 to 142-145, the hydrophobic “anvil-like head” motif (stretch of residues 112-134/141) that is proximal to the C-terminal flank and more than half of the middle β-arch hairpin (residues 125-168) (**S7 Fig**). According to the structure of 263K asymmetric unit, it exists multiple interactions between PrP^res^ N-terminal (region 95-150) and C-terminal flanks (region 155-176), which should contribute to stabilize the 263K filament (**S7 Fig**). Those interactions should not exist in LMW PrP^res^ assemblies and should affect the overall stability (**S8 Table**). It will be of upmost interest to see whether LMW C-terminal flanks would keep a similar architecture alone. It must be noted that the purified PrP^res^ preparations used for cryo-EM were heterogeneous, with presence of spherical objects, suggesting that different structures could coexist. One may be more compatible with LMW PrP^res^.

## Materials and Methods

### Biosafety

Formation of rPrP assemblies, prion protein biochemistry, mouse experimentation and histopathology were performed in biosafety level 2 or level 3 facilities with strict adherence to safety protocols. Atomic force microscopy analyses were performed in biosafety level 1 laboratory with dedicated sample holders and equipment according to approved biosafety procedures. All safety procedures were reviewed and approved by Local and National Prion Research Committees.

### Formation of recombinant PrP fibrillar assemblies

Full-length rPrP with hamster, human and mouse amino acid sequence were produced in *Escherichia coli* and purified as previously described [75]. Purified, monomeric forms of rPrP were stored lyophilized. Conversion of the monomers into amyloid like assemblies was performed as previously described [26, 76]. Briefly, the lyophilized proteins were dissolved directly in 50 mM MES buffer, pH 6.0 (1 mg/mL final concentration). This solution was kept on ice before starting the experiment. To prepare a 600 μL reaction, the following reagents were mixed in a conical plastic tube: water (90 μL), GdnHCI (6 M, 200 μL), MES buffer (0.5M, pH 6.0, 10 μL) and finally the PrP stock solution (300 μL). The solution was mixed by gentle pipetting to avoid introducing air bubbles. Typically, a 10-mL reaction was made in a 15-mL conical centrifuge tube. The tube (arranged horizontally on the plate surface) was incubated with continuous orbital shaking at 30 rpm (16mm amplitude) at 37 °C. Fibril formation was monitored using a thioflavin T binding assay [75] by diluting aliquots into 10mM Na-acetate buffer, pH 5.0 to a final PrP concentration at 0.3μM and adding thioflavin T to a final concentration of 10μM. Samples were then dialyzed in 10mM sodium acetate, pH 5.0. Fibrils were collected by ultracentrifugation for 45min at 228,147 × *g* using a Beckman Optima TL100 ultracentrifuge and a TLA-100.3 rotor, and resuspended in 10 mM sodium acetate, pH 5.0. A washing step was performed by repeating the ultracentrifugation and resuspension steps. All concentrations given for fibrillar PrP refer to the respective equivalent monomer concentration.

### Formation of recombinant PrP oligomers

O1 oligomers were generated as previously described [40]. Sheep rPrP (V_136_R_154_Q_171_ allele) was recovered in 20mM sodium citrate buffer pH 4.1 at 50 μM final protein concentration and incubated at 55°C for 6 h. The product of this incubation was then loaded on TSK 4000SW column equilibrated with the same buffer. The first peak of the elution, which corresponds to O1, was sampled and directly used for inoculation.

### Atomic force microscopy and dynamic light scattering

AFM experiments were performed on a NanoWizard 3 (JPK) in liquid, using a nanoprobe of 10Å diameter (Peakforce-HIRS-F-B2) in a liquid bio-cell chamber in a QI mode. Recombinant PrP assemblies at 0.5mM (concentration expressed in monomer equivalent) were deposited on a freshly cleaned mica surface for 3 minutes. After two iteratives washes of the mica surface with sodium acetate buffer (10mM, pH5.0), the bio-cell compartment was fulfilled with 1mL buffer. The size of the different types of assemblies was estimated using a combination of JPK image analysis interface and a homemade MATLAB code. The mean average hydrodynamic size of the assemblies has been determined by dynamic light scattering.

### Animal experiments

All the experiments involving animals were carried out in strict accordance with EU directive 2010/63 and were approved by INRAE Local Ethics Committee (Comethea; permit numbers 12-034, 15-056 and APAFIS#29603-2021020914525215). The hamster PrP tg7 line has previously been described [3, 27, 77]. This line is homozygous for the hamster PrP transgene and has been established on the Zurich I PrP^0/0^. Level of overexpression is approx. 4-fold compared to golden Syrian hamster brain (**S2 Fig**). These mice do not develop any abnormal phenotype or neurological signs with aging and have a normal life span around 1.5-2 years. They do not develop any spontaneous prion disease upon inoculation with uninfected brain material. Only tg7 females were used; they were 6-8-week-old at the time of inoculation. All mice were group housed by 4 to 5 in polypropylene cages in a standard temperature- and humidity-controlled biosafety laboratory 3 animal facility with a 12-hour light-dark rhythm, unlimited access to food and water and enrichment (igloos, wood toys, nests). Cages, food, enrichment and water were sterilized before use.

### Mouse bioassays

To avoid any cross-contamination, a strict protocol was followed, based on the use of disposable equipment and preparation of all inocula in a class II microbiological cabinet. rPrP assemblies, oligomers and monomers were prepared at the 0.5 mg/mL concentration in 5% w/v glucose extemporarily. Groups of individually identified tg7 mice (8-10 mice per group) were intracerebrally inoculated with 20 μL of the solution, using a 27-gauge disposable syringe needle inserted into the right parietal lobe. Animals were anesthetized with 3% isoflurane during the procedure and placed on a heating pad until they fully recovered. They were monitored daily for general health. They were euthanized at terminal stage or at end-life by cervical column disruption. Their brains were carefully collected with separate, disposable tools, homogenized at 20% w/v in 5% glucose and stored at −80°C until further use. For subpassaging, 20 μL of the solution were intracerebrally reinoculated at 10% w/v to groups of individually identified tg7 mice (5-8 mice per group). The procedure was the same as above. Animals at terminal stage of disease or at end-life were euthanized by cervical column disruption. At each passage, brains were also directly frozen on dry ice (for histoblotting) before storage at −80°C or fixed by immersion in neutral-buffered 10% formalin (for lesion profiling).

### Analysis of PK-resistant PrP^Sc^ by immunoblot

PrP^res^ was extracted from 20% brain homogenates with the Bio-Rad TeSeE detection kit, as previously described [45, 78]. Briefly, 200 μL aliquots were digested with proteinase K (200 μg/mL final concentration in buffer A) for 10 min at 37°C before precipitation with buffer B and centrifugation at 28,000 × *g* for 5 min. Pellets were resuspended in Laemmli sample buffer, denatured, run on 12% Bis-Tris Criterion gels (Bio-Rad, Marne la Vallée, France), electrotransferred onto nitrocellulose membranes with primary and secondary antibodies, as described in **S9 Table**. Immunoreactivity was visualized by chemiluminescence (Pierce ECL, Thermo Scientific, Montigny le Bretonneux, France). Chemiluminescent signals were acquired with the Chemidoc digital imager and analyzed with Image Lab software (Bio-Rad, Marne la Vallée, France). Enzymatic deglycosylation was performed on denatured PrP^res^ with 1,000 U of recombinant PNGase (peptide N-glycosidase F; New England Biolabs) for 2h at 37°C in 1% Nonidet P40 and the manufacturer’s buffer.

### Analysis of thermolysin-resistant PrP^Sc^ by immunoblot

Mouse brain homogenates were diluted in lysis buffer (2% sodium deoxycholate, 2% Triton X-100, 200 mM Tris-HCl pH7.4) containing thermolysin (final concentration 125 μg/mL) and incubated for 1h at 70°C under constant agitation at 800 rpm. The samples were analyzed by electrophoresis and immunoblotted as above.

### PrP^Sc^ insolubility assay

The entire procedure was performed at 4°C. Aliquots (6 μL) of tg7-brain homogenates were supplemented with a mixture of protease inhibitors (Roche), mixed with 5% glucose (94 μL) before adding an equal volume of solubilization buffer (50 mM HEPES pH 7.4, 300 mM NaCl, 10 mM EDTA, 4% w/v dodecyl-β-D-maltoside) and incubation for 45 min. Sarkosyl (N-lauryl sarcosine; Sigma) was added to a final concentration of 2% w/v and the incubation continued for a further 45 min, as described in [3]. The mixture was then loaded on a 40% w/v sucrose gradient before centrifugation at 100 000 x *g* for 1 h at 4°C, using a Beckman Optima TL100 ultracentrifuge. This generated supernatant and pellet fractions. Proteins in the pellet were resuspended in Laemmli sample buffer, denatured and analysed by western blot, as described above.

### Histoblots

Brain cryosections were cut at 8-10 μm (NX-70, MM, Lyon, France), transferred onto Superfrost slides and kept at −20°C until use. Histoblot analyses were performed on 2-3 brains per passage, as previously described, using the 3F4 anti-PrP antibody (human PrP epitope at residues 109 to 112 [31]) or SAF84 anti-PrP antibody (human PrP epitope at residues 160-164 [32]). Analysis was performed with a digital camera (Coolsnap, Photometrics, Paris, France) mounted on a binocular glass (SZX12, Olympus, Paris, France).

### Vacuolar lesion profiles

Haematoxylin-eosin-stained paraffin-embedded brain tissue sections were used to establish standardized vacuolar lesion profiles in mice, as previously described [38, 79]. Analyses were performed on 3-5 brains per passage.

### Statistical analysis

GraphPad Prism 9.0 software (GraphPad, La Jolla, CA, USA) was used to establish the Kaplan-Meier curves plotting the percentage of mice without prion disease against the incubation time. This software was also used to draw the vacuolar profiles.

## Acknowledgments

We thank the staff of Animalerie Rongeurs (INRAE, Infectiology of fishes and rodent facility, doi: 10.15454/1.5572427140471238E12, Jouy-en-Josas, France) for animal care. We thank Byron Caughey (NIH, RML, Montana, USA), Jan Langeveld (Wageniggen University, NL) and Stéphanie Simon (CEA Sacaly, France) for providing us with R20, 12B2 and SAF34 antibodies, respectively.

---

**S1 Fig.** Flow diagram describing the protocol used to prepare recombinant PrP assemblies

**S2 Fig.** PrP^C^ overexpression in hamster PrP transgenic mice does not impact the magnitude of the species barrier for BSE prion strains

**S3 Table.** Absence of prion disease in tg7 mice after intracerebral inoculation of monomers from hamster, mouse and human recombinant PrP

**S4 Fig.** Distinct batches of hamster and mouse recombinant PrP assemblies are pathogenic in tg7 mice

**S5 Fig.** Neuropathological pattern of PrP^res^ deposition in tg7 mice inoculated with recombinant PrP assemblies

**S6 Text.** Estimation of the relative infectious titer of rPrP^Ha^ assemblies

**S7 Fig.** Localization of the two LMW PrP^res^ species on the 3D structure of 263K and RML prions

**S8 Table.** The energetic cost of the deletions 95-145 and 95-155 based on the atomic structure of 263K

**S9 Table.** List of antibodies used in the study

