## Supporting information for "Pathogenicity, strain properties and interspecies transmission capacity of pure recombinant prion protein assemblies"

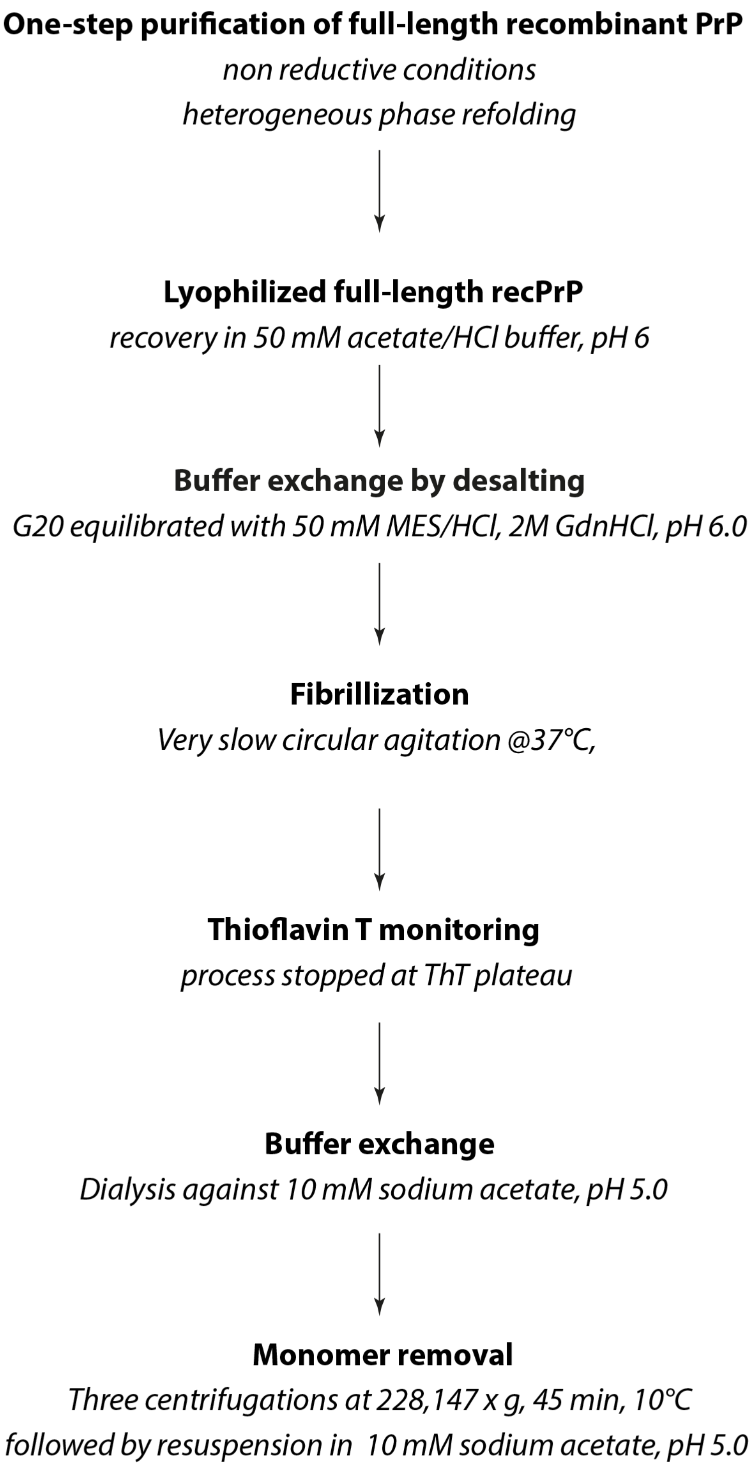


**S1 Fig. Flow diagram describing the protocol used to prepare recombinant PrP assemblies**

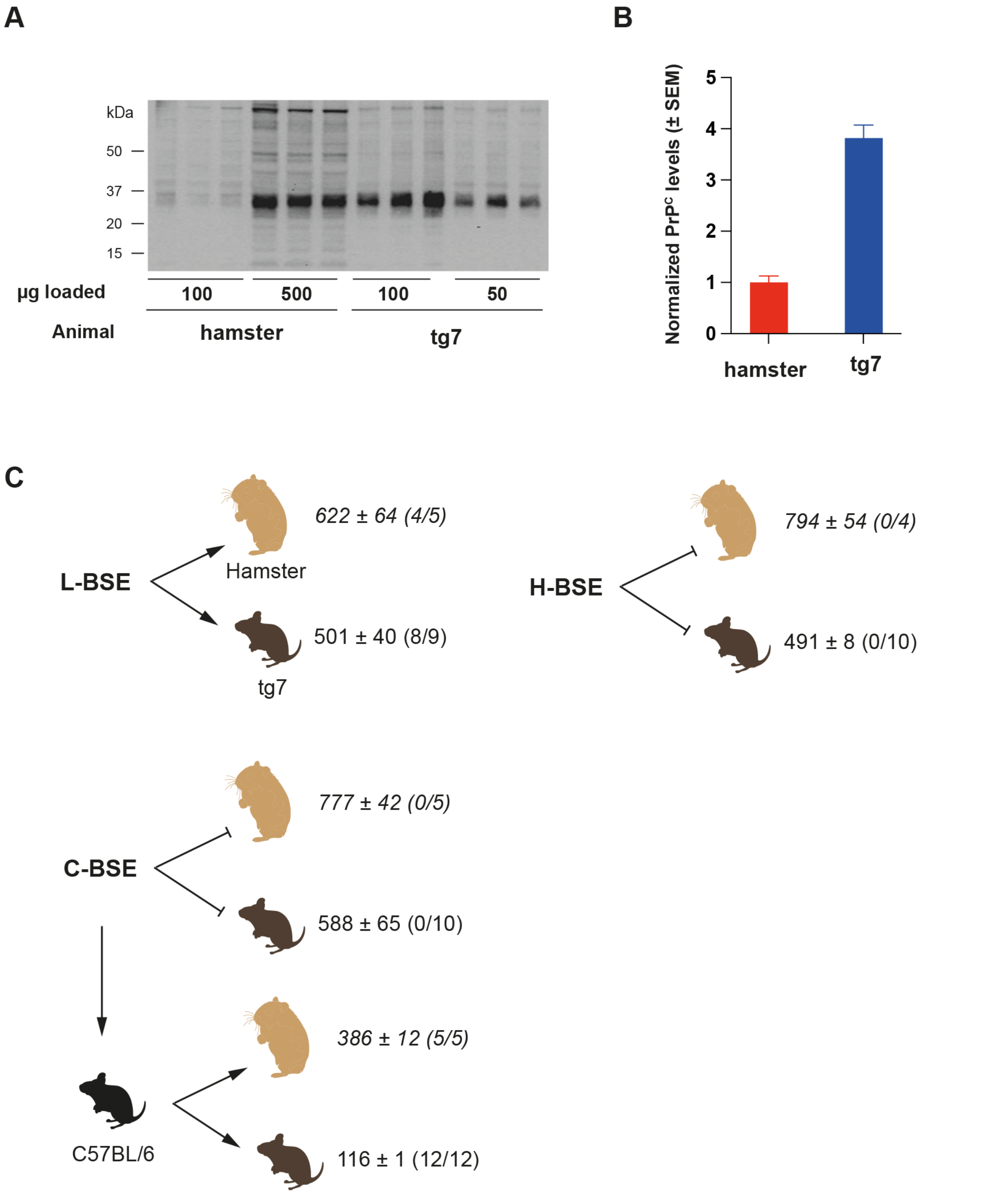


**S2 Fig PrP$^{C}$ overexpression in hamster PrP transgenic mice does not impact the magnitude of the species barrier for BSE prion strains**

(**A**) Immunoblot (Sha31 antibody) and (**B**) quantification of $PrP^C$ overexpression in the brain of hamster PrP mice (tg7) as compared to that in the brain of golden Syrian hamster. Quantification was performed on the immunoblots on *n*=3 animals. Results are expressed as mean levels ± SEM. (**C**) Comparison of the magnitude of the species barrier on transmission of BSE prions to golden Syrian hamsters or to hamster PrP transgenic mice. Atypical BSE prions (L-BSE, H-BSE), classical BSE prions (C-BSE) before or after intermediate passage in C57BL/6 mice were transmitted by intracranial route to tg7 transgenic mice expressing hamster PrP (protocol as previously described [1, 2]). For comparison, the transmission properties of the same prion strains in golden Syrian hamster are shown (data in italics from [3]). The mean survival time±SEM (in days) are indicated. The number of disease and/or $PrP^{Sc}$-positive animals / number of inoculated animals are in bracket. The force of species barrier is similar between hamsters and transgenic mice expressing hamster PrP.

**S3 Table. Absence of prion disease in tg7 mice after intracerebral inoculation of monomers from hamster, mouse and human recombinant PrP**

| | Range of individual survival time in days ($n/n_0$)[a] | | |
|---|---|---|---|
| PrP monomers | Hamster | Mouse | Human |
| 1st passage | 488-718 (0/8) | 464-724 (0/8) | 456-660 (0/11) |

[a]$n/n_0$: number of mice with neurological disease and positive for PrP$^{res}$ in the brain by immunoblotting/number of inoculated mice

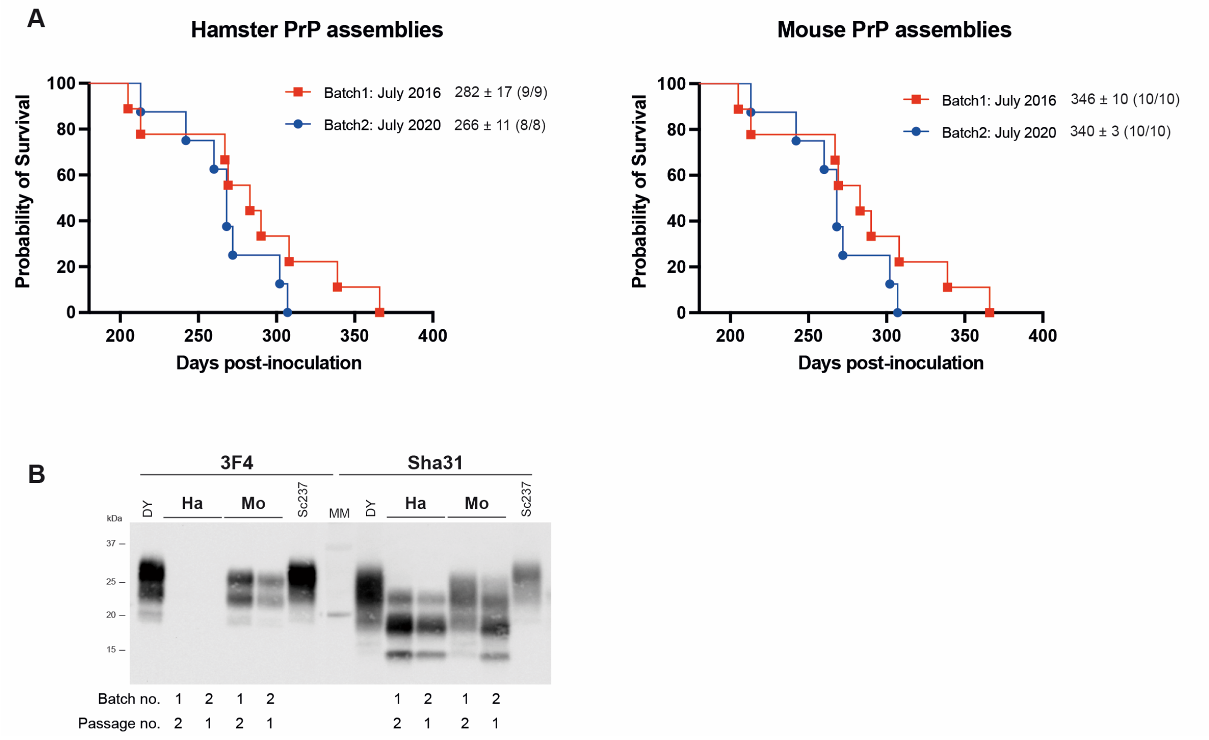


**S4 Fig. Distinct batches of hamster and mouse recombinant PrP assemblies are pathogenic in tg7 mice**

(**A)** Kaplan-Meier curve plotting the percentage of mice without prion disease (survival) against the incubation time (days post-inoculation). The different colors and symbols describe the inoculated batches. The hamster and mouse batches were prepared independently with a 4-year interval. The survival curves for batch 1 and batch 2 were not statistically significantly different (logrank test). The survival expressed as mean ± SEM days and in parenthesis the number of diseased, $PrP^{res}$-positive/number of inoculated mice are indicated for each batch.

(**B**) Electrophoretic pattern of $PrP^{res}$ in the brains of tg7 mice inoculated with different batch of assemblies from hamster (Ha) or mouse (Mo) rPrP. The 3F4 and Sha31 monoclonal anti-PrP antibodies were used to probe the blots, to detect HMW and HMW+LMW $PrP^{res}$, respectively. Tg7-passaged Sc237 and DY $PrP^{res}$ are shown as control. MM: molecular mass markers.

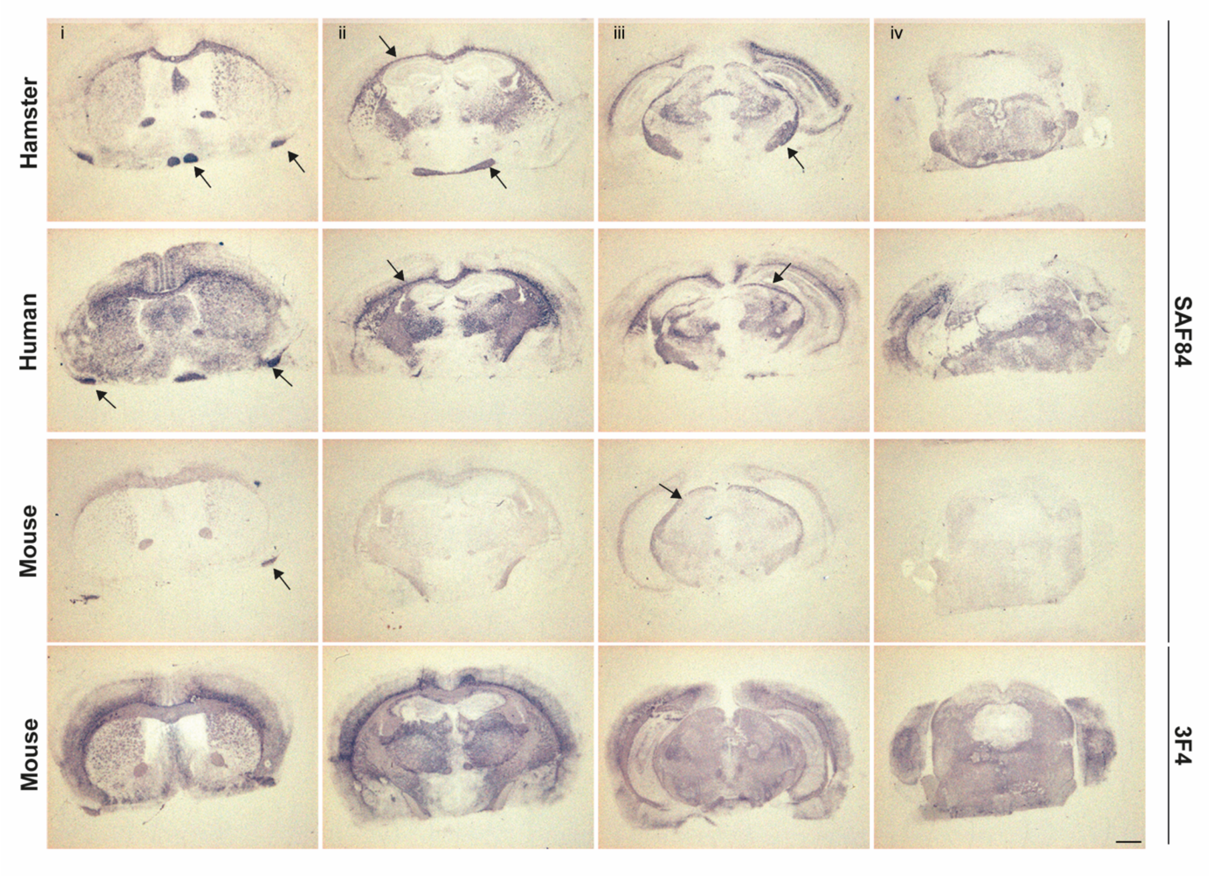


**S5 Fig. Neuropathological pattern of PrP$^{res}$ deposition in tg7 mice inoculated with recombinant PrP assemblies**

Representative histoblots of rostro-caudal transversal brain sections are shown at the 3$^{rd}$ passage, after challenge with rPrP assemblies from hamster, human and mouse species. Analyses were performed at the level of the septum (i), hippocampus (ii), midbrain (iii) and brainstem (iv). Histoblots were probed with 3F4 and SAF84 anti-PrP monoclonal antibody, as indicated. Arrows indicate PrP$^{res}$ deposition in white matter tracts. Scale bar, 1mm.

**S6 Text. Estimation of the relative infectious titer of $rPrP^{Ha}$ assemblies**

We inoculated 10 µg of $rPrP^{Ha}$ assemblies at primary passage, i.e. a concentration of 400 pmoles. Comparing this value to that of $PrP^{Sc}$ in a prion-containing brain material remains difficult. It has been reported that 263K-infected hamster brain (at terminal stage of disease) may contain ≈20 µg of $PrP^{Sc}$ / g of brain, i.e. a $C_{tot}$ of ≈$8.10^{-10}$ mol/g brain [4-6]. Tg7 mice accumulate 5-10 less $PrP^{Sc}$ than hamster, thus $C_{tot}$ may be ≈$1.10^{-10}$ mol/g brain. As can be seen in **Fig 2A** (panel + PNGase), tg7 mice accumulate ≈1.4-fold more $PrP^{Sc}$ after $rPrP^{Ha}$ inoculation than after Sc237/263K inoculation Thus $C_{tot}$ may be ≈$1.4.10^{-10}$ mol/g brain. As 2 mg of brain are inoculated, this means that ≈28 pmoles of $PrP^{Sc}$ are injected on second passage of $rPrP^{Ha}$-derived prions. Thus, there would be a 15-fold difference in terms of PrP species injected between the 1st and the 2nd passage for $rPrP^{Ha}$, which would be consistent with the smooth reduction of incubation time over passaging and suggests relatively high infectivity levels of the $rPrP^{Ha}$ preparations on primary passage.

Compared to tg7-passaged 263K/Sc237, there would be a 20-fold difference in terms of PrP species injected at primary passage of $rPrP^{Ha}$. This estimation suggests that the concentration of recombinant PrP assemblies inoculated at primary passage is not aberrantly high.

**263K prions**

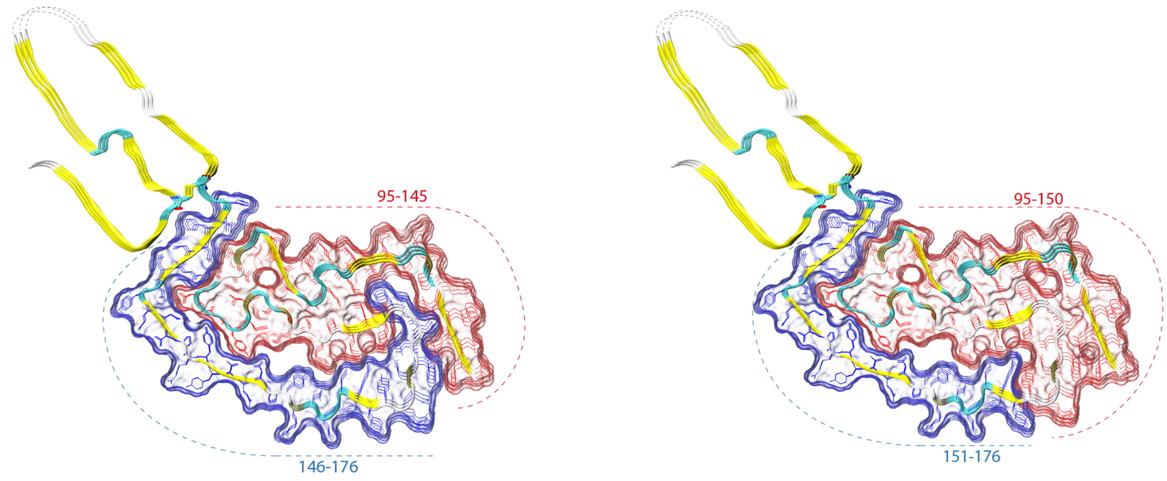


**RML prions**

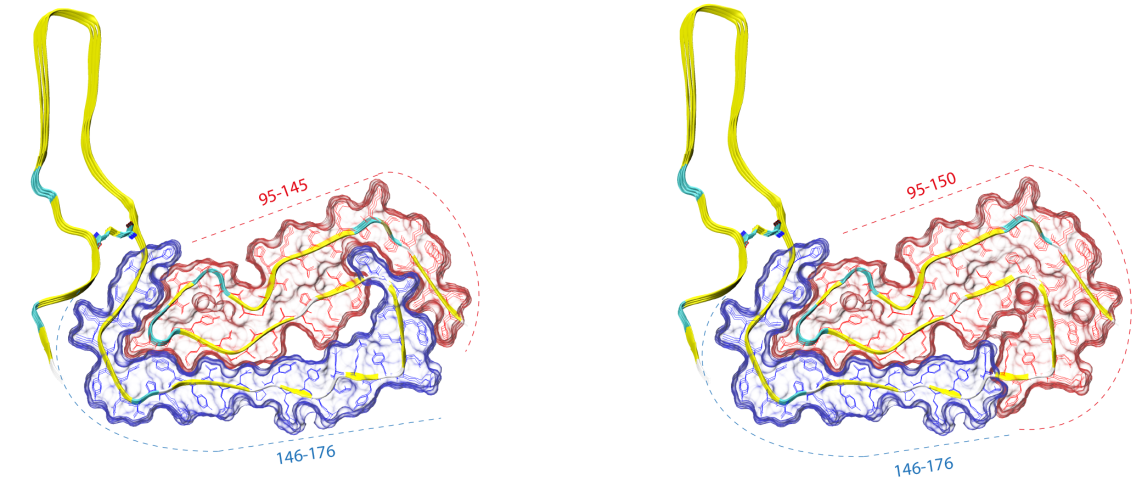


**S7 Fig. Localization of the two LMW PrP$^{res}$ species on the 3D structure of 263K and RML prions**

The LMW species generated during the replication of the synthetic prions comes from the conversion of full-length PrP$^{C}$ and endogenous truncation or from conversion of a truncated form of PrP$^{C}$. Epitope mapping experiments showed that the two LMW species are truncated at residues around positions 145 and 150. By assuming that the variants of PIRBS-fold constitute a generic structure for prion assemblies, we analyzed the energetic cost of regions 95-145 or 95-150 deletion on the stability of the asymmetric unit by using as model the structure of 263K and RML asymmetric unit (PDB code respectively 7LNA and 7TD6). For our estimation we used FoldX force field [7]. As can be seen in **S8 Table A**, the domain 95-145 contributes approximatively to 60% of the asymmetric unit stability. The contribution of 95-150 to the global stability is in the same order of magnitude. Furthermore, the region 95-145 appears to play a significant role in the stacking stabilization (**S8 Table B**).

**S8 Table. The energetic cost of the deletions 95-145 and 95-155 based on the atomic structure of 263K**

**(A)** The effect of the deletion 95-145 and 95-155 on the global stability of the 263K asymmetric unit as estimated using FoldX software. The values are in kcal/mol.

| | 263K | D95-145 | D95-155 |
|---|---|---|---|
| BackHbond | -203.89 | -116.48 | -108.66 |
| SideHbond | -112.2 | -19.64 | -19.64 |
| Energy_VdW | -402.1 | -250.45 | -226.45 |
| Electro | -7.52 | +4.98 | +6.71 |
| Energy_SolvPolar | +541.66 | +376.92 | +342.28 |
| Energy_SolvApolar | -506 | -305.34 | -276.90 |
| Energy_vdwclash | +25.76 | +48.05 | +45.16 |
| energy_torsion | +14.42 | +22.30 | +21.73 |
| backbone_vdwclash | +67.46 | +39.03 | +35.00 |
| **Total** | **-582.41** | **-200.63** | **-188.77** |

**(B)** The contribution of the region 95-145 on the protomer stacking stabilization as estimated by performing the deletion 95-145 either on chain A or chain B of the asymmetric unit. The values are in kcal/mol.

| | 263K | Dchain A 95-145 | Dchain B 95-145 |
|---|---|---|---|
| BackHbond | -203.89 | -112.91 | -149.22 |
| SideHbond | -112.2 | -22.09 | -25.09 |
| Energy_VdW | -402.1 | -265.53 | -321.25 |
| Electro | -7.52 | +6.89 | +12.92 |
| Energy_SolvPolar | +541.66 | +399.26 | +476.98 |
| Energy_SolvApolar | -506 | -327.88 | -395.73 |
| Energy_vdwclash | +25.76 | +54.58 | +57.05 |
| energy_torsion | +14.42 | +24.87 | +25.49 |
| backbone_vdwclash | +67.46 | +49.34 | +53.00 |
| **Total** | **-582.41** | **-193.47** | **-265.85** |

**S9 Table. List of antibodies used in the study**

| Anti-PrP antibody | | Supplier | Reference |
|---|---|---|---|
| Sha31 | Mouse IgG1 | Cayman Chemical | [8] |
| 3F4 | Mouse IgG2a | Merck | [9] |
| 12B2 | Mouse IgG1 | Jan Langeveld | [10] |
| SAF34 | Mouse IgG | Stéphanie Simon | [8] |
| mAb132 | Mouse IgG1 | Creative Biolabs | [11] |
| SAF84 | Mouse IgG | Cayman | [8] |
| R20 | Rabbit polyclonal | Byron Caughey | [11] |
| EP1802Y | Rabbit monoclonal | Abcam | [11] |
